# The G256E HCM mutation prolongs relaxation via altered nucleotide handling

**DOI:** 10.1101/2025.08.29.673178

**Authors:** Kerry Y. Kao, Matthew Carter Childers, Divya Pathak, Rama Reddy Goluguri, Timothy S. McMillen, Kathleen M. Ruppel, James A. Spudich, Michael Regnier

## Abstract

Mutations in myosin alter its motor functions in diverse ways by affecting different structural and chemo-mechanical events. Multidisciplinary strategies can be used to understand how varying alterations in motor function converge to common phenotypes like hypercontractility and hypertrophic cardiomyopathy (HCM). Here, we combined molecular dynamics (MD) simulations with protein biochemical and myofibril mechanical analyses to study the HCM-causing myosin variant G256E. MD simulations demonstrated that G256E induces structural changes that increase the work required to displace ADP.Mg^2+^ from actomysoin complex. Stopped-flow biochemical analysis demonstrated increased ADP affinity for actomyosin and single myofibril mechanics analysis demonstrated increased force generation and reduced ADP sensitivity of the early, slow phase of relaxation. Together, these results demonstrate that slower ADP release from myosin during contraction is a significant contributor to pathological contractile nature of the G256E mutation. This study highlights the importance of detailed chemo-mechanical analysis of mutations associated with hereditary cardiac diseases.

## 2. Introduction

Hypertrophic cardiomyopathy (HCM) affects 1 in 500 people globally and is implicated as a contributing cause of other life-threatening heart diseases such as atrial fibrillation, heart failure, and arrhythmia resulting in sudden death.^1^ About 60% of HCM cases are caused by familial inheritance of a single autosomal dominant mutation within a gene encoding a sarcomere protein.^2^ In particular, the *MYH7* gene encoding the β-myosin heavy chain (MHC) is one of the two most common hotspots for HCM mutations.^3^ Myosin is the motor protein responsible for muscle contraction, converting chemical energy produced by ATP hydrolysis and release of products (P_i_, ADP) into the mechanical work of force production and muscle cell shortening.^4^ This involves changes in myosin’s structure as it interacts with actin to accelerate the release of ATP hydrolysis products (P_i_, ADP) in a process known as chemo-mechanical transduction.^5–7^ Structural perturbations resulting from disease-associated mutations can disrupt the kinetics and/or mechanics of the chemo-mechanical cycle and can lead to initiation and progression of disease.^8^

The G256E MYH7 mutation, associated with HCM,^9^ is located in a hairpin turn that connects two strands in the central β-sheet of transducer region of the myosin head. This region conveys information between the nucleotide binding pocket and actin interface of myosin, both critical to chemo-mechanical transduction during contraction and relaxation.^10^ In a recent study, we employed CRISPR-Cas9 to edit this mutation into human induced pluripotent stem cell-derived cardiomyocytes (hiPSC-CMs) to produce a heterozygous (+/-) isogenic model for studying its impact at multiple scales of cardiac muscle organization.^8^ A biochemical single ATP turnover assay performed on purified human β-cardiac 25-hep heavy meromyosin (HMM) containing the G256E mutation results in a reduced population of myosin in a low basal ATPase state, often described as the super-relaxed (SRX) state, suggesting a greater portion of the myosin population is available to contribute to muscle contraction. Comparison of the actin-activated ATPase activity of two versions of G256E myosin that differed only in the length of their coiled-coil tail confirmed that the mutation disrupted the ability of myosin to fold back into a sequestered, low ATPase state. This supports the hypothesis that the mutation results in more heads being available to interact with actin at the beginning of contractions. Slower motility in an *in* vitro F-actin motility assay indicated the mutation results in a higher myosin duty ratio, suggesting that the actomyosin detachment rate may also be affected. This idea was supported by measures of the contraction and relaxation properties of isolated myofibrils from hiPSC-CMs. The kinetics of the slow, early phase of relaxation, which measures the myosin detachment rate from thin filaments,^11^ was slower for G256E versus WT myofibrils. Molecular dynamics (MD) simulations of the myosin M.ATP-state suggested that G256E disrupts the shape and stability of the central β-sheet, potentially altering communication between the nucleotide-binding pocket and actin-binding surface. This communication path is critical to open the nucleotide binding cleft that allows ADP release. This, in turn, is required before a new ATP molecule can bind in the nucleotide pocket and lead to the dissociation of myosin from actin. While these findings are suggestive that myosin-actin chemo-mechanics are altered as part of the hypercontractile phenotype of G256E, it is important to understand which particular step(s) of the cross-bridge cycle are affected.

The ADP release step is rate limiting for cross-bridge formation in sarcomeres during loaded contractions and is also a determinant of the rate of relaxation.^12^ This is supported by biochemical,^13^ mechanical,^14^ single molecule,^15^ and structural^16^ evidence, and holds true across multiple classes of myosins from diverse species. Recent studies have shown that various myosin mutations can alter the affinity of myosin for ADP and consequently affect cross-bridge cycling kinetics.^17^ Thus, detailed study of how the G256E mutation affects ADP release during chemo-mechanical cycling is merited.

Here we report the effects of myosin G256E on ADP release from the nucleotide binding pocket of myosin and its effects on myosin and myofibril chemo-mechanical function. MD simulations of post-powerstroke myosin containing ADP (with Mg^2+^) complexed with actin (AM.ADP) suggested that G256E significantly alters nucleotide pocket dynamics. Steered MD simulations suggested that the sequence and timing of contact pair disruptions in the pocket are altered and that more energy is required to pull ADP from the pocket. These results provide a structural basis for altered ADP-myosin affinity and dynamics. Stopped-flow kinetics analysis showed that G256E sS1 had a ∼30% increase in ADP affinity over wild type (WT) sS1, validating the MD simulation predictions.

To determine how this affects the chemo-mechanical function of myofibrils, we measured contraction and relaxation in the absence and presence of elevated ADP. Similar to our initial study,^8^ G256E myofibrils generated significantly more force and exhibited a slower, early linear phase kinetics of relaxation (*k*_REL,slow_) compared to isogenic controls. When ADP was elevated in activating and relaxation solutions there was considerably less effect on *k*_REL,slow_ for G256E myofibrils, suggesting that ADP release is impaired with the mutation.

Combined, these experimental and computational studies support the hypothesis that the G256E mutation alters the structure of the nucleotide binding pocket, resulting in greater ADP affinity for G256E myosin than WT myosin, and this results in slower ADP release to slow crossbridge detachment and prolong contractile relaxation. To our knowledge, this is the first study to examine the reaction pathway and free energy changes associated with release of Mg^2+^-ADP from a post-powerstroke-like state of the actomyosin complex at the atomic scale with molecular simulations. These results highlight the power of combining experimental measures with in-depth computational dynamics to provide detailed structure-based mechanistic insights at the atomistic scale for protein and contractile organelle (myofibril) function.

## 3. Materials and Methods

### 3.1 Computational Model Preparation

The initial actin+myosin+ADP+Mg^2+^ (AM.ADP-Mg^2+^) structure model was generated based on cryo-EM structures of post-powerstroke actin-bound human cardiac myosin solved by Doran *et al.* (PDB ID: 8EFH, 3.3 Å resolution; PDB ID: 8EFE, 3.8 Å resolution).^16,18^ Our model includes the myosin motor and actin 5-mer from 8EFH; tropomyosin was excluded (chains O, P). Then we grafted the converter domain, tail and essential light chain (ELC) from 8EFE onto the actomyosin complex. Missing disordered heavy atoms were constructed via *Modeller*^19^. The disordered N-terminal extension of the ELC was not modeled. The final WT model includes actin residues (2-377), myosin residues (1-810), and ELC residues (44-195). A model for the mutant G256E actomyosin complex was generated via *in silico* mutation of the WT structure with *UCSF Chimera.*^20,21^

The resulting two systems (WT and G256E) were prepared for molecular dynamics (MD) simulation using the AMBER20 simulation package^22^ and the ff14SB force field^23^. Water molecules were treated with the TIP3P force field.^24^ Metal ions were modeled using the Li and Merz parameter set.^25–27^ ADP molecules were treated with the GAFF2 force field.^28^ Partial charges for ADP were derived from a restrained electrostatic potential (*resp*) fit to quantum mechanics calculations performed with ORCA.^29^ The SHAKE algorithm was used to constrain the motion of hydrogen-containing bonds.^30^ Long-range electrostatic interactions were calculated using the particle mesh Ewald (PME) method. Hydrogen atoms were modeled onto the initial structure using the *leap* module of AMBER, and each protein was solvated with explicit water molecules in a truncated octahedral box and then neutralizing + 120 mM counterions (Na^+^, Cl^-^) were added. Each system was minimized in three stages. First, hydrogen atoms were minimized for 1000 steps in the presence of 100 kcal/mol restraints on all heavy atoms. Second, all solvent atoms were minimized for 1,000 steps in the presence of 25 kcal/mol restraints on all protein atoms. Third, all atoms were minimized for 8,000 steps in the presence of 25 kcal/mol restraints on all backbone heavy atoms (N, O, C, and C_α_ atoms). After minimization, systems were heated to 310 K using the NVT (constant number of particles, volume, and temperature) ensemble and in the presence of 25 kcal/mol restraints on backbone heavy atoms. Next, the systems were equilibrated over five successive stages using the NPT (constant number of particles, pressure, and temperature) ensemble. During the first four stages, the systems were equilibrated for 5 ns in the presence of 25 to 1 kcal/mol restraints on backbone heavy atoms. During the final equilibration stage, the systems were equilibrated for 5 ns in the absence of restraints. A 10 Å nonbonded cutoff was used for all preparation and production simulations.

### 3.2 Conventional MD Production and Analysis

The equilibrated systems were then simulated using conventional molecular dynamics protocols in the NVT ensemble in triplicate for 500 ns each (2 systems, three replicate simulations per system, 500 ns sampling per replicate simulation = 3 µs net sampling) and coordinates were saved every 10 ps.

Molecular analyses were performed using *cpptraj*.^31^ C RMSD values for each simulation were calculated for all residues and relative to the minimized starting structure. C_α_ RMSF values were calculated after aligning the trajectory to the average structure. Residue-residue contacts were counted if at least one pair of heavy atoms between two residues were within 5 Å of one another and contacts are presented as the fraction of simulation time that two residues were interacting. Nucleotide dynamics were evaluated by monitoring 6 dihedral angles that describe ADP structure using internal coordinates, as previously described.^32^ The 6 calculated dihedral angles were defined as: θ_1_ (atoms: O1B-PB-O3A-PA), θ_2_ (PB-O3A-PA-O5′), θ_3_ (O3A-PA-O5′-C5′), θ_4_ (PA-O5′-C5′-C4′), θ_5_ (O5′-C5′-C4′-O4′), and θ_6_ (O4′-C1′-N9-C8). Ramachandran-style dihedral angle maps (θ_1_ vs θ_2_, θ_3_ vs θ_4_, and θ_5_ vs θ_6_) describe the relative orientation of the phosphate groups, the sugar ring, and the purine ring. The orientation of the myosin tail relative to the actin filament was calculated using the ‘vector’ module of *cpptraj* as previously described.^33^ We also calculated interaction energies between myosin residues and either ADP or Mg^2+^ using the linear interaction energy module. Electrostatic and van der Waals interaction energies were combined.

### 3.3 Steered molecular dynamics (sMD)

Pre-production protocols were identical to those for the conventional MD simulations except that we employed the hydrogen mass partitioning scheme^34^ to enable a 4 fs time step, the total time for the heating and equilibration phases was 1 ns, and 25 kcal/mol restraints were present on all protein heavy backbone atoms, ADP, and Mg^2+^ during the entire equilibration. During the production simulations we steered ADP.Mg^2+^ 14.4 Å away from the nucleotide binding pockets using a center of mass distance-based reaction coordinate. ADP.Mg^2+^ was steered away at 1 Å/nanosecond and so production simulations were 10.8 ns long (WT) or 10 ns long (G256E). 60 WT and 60 G256E simulations were performed. We calculated the average work done to displace ADP, the free energy profiles along the reaction coordinate, the Jarzynski average trajectories, and the average myosin-ADP.Mg^2+^ interactions made along the reaction coordinate.

### 3.4 sS1 protein purification

Recombinant WT and G256E human β-cardiac sS1 myosins were purified as described previously.^35^ Briefly, human β-cardiac myosin heavy chain (*MYH7)* was co-expressed with human ELC (*MYL3)* in the differentiated murine myoblast C2C12 cell line (ATCC) using adenovirus generated in HEK293T cells (ATCC) using the AdEasy Vector System (Qbiogene Inc, USA). The sS1 construct is a truncated version of MYH7 (residues 1-808) followed by a flexible GSG (Gly-Ser-Gly) linker and a carboxy-terminal 8-residue (RGSIDTWV) PDZ binding peptide. The MYL3 construct contains an N-terminal FLAG tag followed by a TEV protease site for purification. C2C12 cells were infected with adenovirus constructs 48 hr after differentiation and harvested 4 days after infection in a lysis buffer containing 50 mM NaCl, 20 mM MgCl_2_, 20 mM imidazole pH 7.5, 1 mM EDTA, 1 mM EGTA, 1 mM DTT, 3 mM ATP, 1 mM PMSF, 5% sucrose, and Roche protease inhibitors. The harvested cells were immediately flash frozen in liquid nitrogen. The cell pellets were stored up to 2 months at –80°C and thawed at room temperature for purification. Cells were lysed in a dounce homogenizer with 50 strokes and the lysate was clarified by spinning at 30,000rpm in a Ti-60 fixed angle ultracentrifuge rotor for 30 min at 4°C. Supernatant was transferred to a fresh tube and bound to anti-FLAG resin for 1.5hr at 4°C on a nutator. The resin was washed with a buffer (150 mM NaCl, 5 mM MgCl_2_, 20 mM imidazole pH 7.5, 1 mM EDTA, 1 mM EGTA, 1 mM DTT, 3 mM ATP, 1 mM PMSF, 10% sucrose, and Roche protease inhibitors) and then nutated overnight with TEV protease at 4°C to cleave the ELC-myosin complex from the resin. The next day, the supernatant was purified using a HiTrap Q HP column on a fast protein liquid chromatography (FPLC) apparatus with a gradient of 0–600 mM NaCl over 20 column volumes in a buffer containing 10 mM imidazole pH 7.5, 4 mM MgCl_2_, 10% sucrose, 1 mM DTT, and 2 mM ATP. Fractions containing human β-cardiac myosin and devoid of mouse skeletal myosin (Coomassie staining on 12.5% SDS PAGE) were pooled and flash frozen in liquid nitrogen. Purified sS1 was buffer exchanged into stopped-flow buffer (120 mM KCl, 5 mM MgCl_2_, 20 mM MOPS, 1 mM DTT, pH 7.0) overnight prior to experiments. Concentration was measured through a Bradford protein assay (5000201, Bio-Rad).

### 3.5 Pyrene-actin labeling

Pyrene-actin was prepared fresh and used within ∼1 week. First, G-actin was prepared from lyophilized bovine cardiac muscle as described before.^36^ G-actin was diluted to 1 mg/mL before polymerizing with 1 mM ATP, 50 mM KCl, and 2 mM MgCl_2_ at 4 °C for 2 hrs. To label, 1.28% w/w N-(1-Pyrene)iodoacetamide (AA00G316, aablocks) was added to F-actin and stirred for 18 hrs at room temperature. Excess N-(1-Pyrene)iodoacetamide was removed by centrifuging at 2,000 x g for 5 mins at 4 °C. Labeled F-actin was pelleted by centrifugation at 92,000 x g for 3 hrs at 4 °C and resuspended in stopped-flow buffer plus minimal ATP (120 mM KCl, 5 mM MgCl_2_, 20 mM MOPS, 1 mM DTT, 200 μM ATP, pH 7.0) in a glass dounce homogenizer. Pyrene-actin was stabilized with 1:1 molar phalloidin (5301, AAT Bioquest) for at least 4 hrs before dialyzing into stopped-flow buffer overnight to remove residual ATP.

### 3.6 Stopped-flow kinetics

Stopped-flow kinetics experiments were performed at 20 °C using a Hi-Tech ‘KinetAsyst’ stopped-flow (SF-61DX2, TgK Scientific, UK). Measurements were made in 120 mM KCl, 5 mM MgCl_2_, 20 mM MOPS, 1 mM DTT, pH 7.0 solution. Pyrene fluorescence was measured by excitation at 365 nm and emission through a GG 400 nm cutoff filter. ADP affinity (*K*_ADP_) was assessed by competitive binding of ATP and ADP to acto-myosin. Briefly, 0.125 μM actomyosin (sS1 + pyrene-actin) was rapidly mixed with 25 µM ATP plus various amounts of ADP (0 – 640 µM). All concentrations reported were post-mixing. An average trace of at least eight individual traces was generated for each condition and fit with a double exponential. The relative *k*_obs_ of the fast phase (*k*_rel_) was plotted vs. ADP concentration and fitted with an equation as described previously.^37^

### 3.7 Human induced pluripotent stem cell (hiPSC) culture

The parental WTC hiPSC cell line was generated by the Bruce R. Conklin Laboratory at the Gladstone Institutes and University of California, San Francisco (UCSF).^38^ Cells were maintained using described methods.^39,40^ A detailed cell culture protocol from the Allen Institute for Cell Science can be found on its webpage (http://allencell.org). AICS-0075-085 ACTN2-mEGFP was developed at the Allen Institute for Cell Science (http://allencell.org/cell-catalog) and is available through Coriell. AICS-0097-113 ACTN2-mEGFP *MYH7*^WT/WT^ and AICS-0097-141 ACTN2-mEGFP *MYH7*^WT/G256E^ were developed at the Allen Institute for Cell Science and are available on request.

### 3.8 hiPSC-Cardiomyocyte (hiPSC-CM) differentiation and maintenance

hiPSCs were differentiated into cardiomyocytes using an established small-molecule protocol^41^ with modifications (Fig. 4c).^42^ Briefly, hiPSCs were seeded onto Matrigel-coated 12-well tissue culture treated plates at densities ranging from 50 Ill 10^3^ – 125 Ill 10^3^ cells per well and maintained for 2 days with mTeSR1 media (85850, Stemcell Technologies). On day -1, the day prior to differentiation, mTeSR1 media was supplemented with 1 μM CHIR99021 (13122, Cayman Chemical). On day 0, hiPSCs were induced with 5 μM CHIR99021 in a chemically defined media composed of RPMI 1640 (11875-093, Gibco) + 500 μg/mL bovine serum albumin (A9418, Sigma-Aldrich) + 213 μg/mL L-ascorbic acid (A8960, Sigma-Aldrich) hereafter abbreviated RBA. After 48 hours, the media was replaced with 2 μM Wnt-C59 (S7037, Selleck Chem) in RBA media. On day 4, the media was replaced with RBA media. From day 6 onwards, differentiated cells were maintained in RPMI 1640 + 1X B27 plus insulin (A1895601, Gibco) with media changes every 2-3 days.

### 3.9 Solutions for myofibril mechanics

Solutions for isolated myofibril mechanics experiments were prepared as previously described.^43^ Experimental solutions were composed of the following: 15 mM total EGTA, 80 mM total MOPS, 5 mM MgATP, 15 mM total creatine phosphate, 83 mM free K, and 52 mM free Na, all free ion concentrations were determined using MAXCHELATOR^44^). Solutions were adjusted to pH 7.0 with NaOH. Elevated ADP solutions (5.6ADP) were made by replacing 50% of the nucleotide pool with ADP for final concentrations of 2.5 mM ATP and 2.5 mM ADP.

### 3.10 Isolated myofibril assay

hiPSCs were differentiated into cardiomyocytes as described above. On day 14 post-differentiation, hiPSC-CMs were replated at 100k cells/cm^2^. On day 16, hiPSC-CMs were subjected to glucose starvation and lactate enrichment for 4 days in DMEM minus glucose (11966-025, Gibco) media supplemented with 4 mM sodium L-lactate (L7022, Sigma Aldrich). hiPSC-CMs were maintained in RPMI 1640 + 1X B27 plus insulin. On day 30, hiPSC-CMs were replated onto 10 kPa polyacrylamide gels in 6-well glass bottom plates (P06-20-1.5-N, Cellvis) patterned with 15 μm width Matrigel lines prepared by microcontact printing as described previously^45^, and matured for 7 days to promote elongation and formation of cell-to-cell contacts. On day 37, isolated myofibrils for mechanical and kinetic analysis hiPSC-CMs were demembranated in a 1% triton-X pCa 9.0 relaxing solution supplemented with 1:100 protease inhibitor cocktail (P8340, Sigma-Aldrich) for 30 minutes on ice. A cell scraper was then used to gently free isolated myofibrils from permeabilized cells. The supernatant was collected and gently homogenized in a glass dounce homogenizer ∼5-7X. Myofibrils were mounted into a custom-built apparatus with rapid Ca^2+^ solution switching (15 °C) as previously described.^43,46^ Myofibrils were initially suspended in pCa 9.0 relaxing solution before rapidly switching to pCa 4.0 maximal activation solution. Myofibrils were also activated in a pCa 5.6 submaximal activation solution and a pCa 5.6ADP solution containing 50% ADP (2.5 mM ATP, 2.5 mM ADP) solution to challenge myofibrils with ADP.

### 3.11 Myosin separation gel

Cell pellets were flash frozen in liquid nitrogen and stored at -80°C until ready to use. Cell pellets were resuspended in ∼3x volume myofibril lysis buffer (5% SDS, 50 mM Tris (pH 8.5), 0.75% sodium deoxycholate, and 1x protease inhibitor cocktail). Sarcomeres and proteins were denatured by boiling samples for 10 minutes. Concentration was measured by blotting for total protein and adjusting loading. Myosin separation gel was made by combining a stacking gel (2.95% Acrylamide with 10% glycerol and pH 8.8) and resolving gel (6% Acrylamide with 10% glycerol and pH 8.8). Separation gels were run for 2 hours and 15 min at 32 mA in a cold room (4°C). Gels were stained for total protein using One-Step Lumitein™ Protein Gel Stain, 1X following manufacturer instructions. Gels were imaged on a BioRad ChemiDoc using SYPRO Ruby protein stain setting.

## 4. Results

### 4.1 G256E alters myosin nucleotide binding pocket dynamics with ADP

To investigate the possible structural changes within myosin that might result in impaired ADP release we employed conventional molecular dynamics (cMD) to simulate the time-dependent changes of the post-powerstroke state of myosin complexed with actin and ADP (AM.ADP, Fig. 1a), and how G256E alters structure and contacts at the atomic level. In general, G256E simulations had a greater change in global C_α_ RMSD compared to WT simulations (Fig. 1b,c), presumably as a response to the change from Gly to Glu. These changes tended to stabilize by 300 ns (Fig. 1c). We calculated the change in C_α_ RMSF per residue and found that the flexibility of the G256E structure remained very similar to WT except for loop 2 (residues 621-646) and the end of the tail (residues 771-810) (Fig. 1d). In the WT stimulations, ADP primarily formed contacts with the P-loop (residues 179-184) and Purine-ring (residues 126-131) of the nucleotide binding pocket (Fig. 2a,c). In G256E simulations, the introduction of the bulky, negatively charged sidechain (E256) formed a novel salt bridge with R169, as was observed previously in simulations of the G256E M.ATP state.^8^ This structural change propagated to the nucleotide pocket. For the majority of G256E simulations, ADP preferred to adopt a conformation that closely associates with Switch-1 (residues 232-244) and the loop that connects the third β-strand of the transducer to the SH1-helix (SH1-loop, residues 673-685). In particular, two non-native contacts formed in the G256E simulations (Fig. 2c S241 & T678) and persisted for greater than 75% of combined simulation time. This new ADP conformation led to the formation of a salt bridge between D239 of Switch-1 and K679 of the SH1-loop, which persisted longer in G256E simulations (average 28.9% of simulation time) compared to WT (average 3.2% of simulation time, p = 0.051). Overall, this results in a loss of contacts between ADP and the P-loop and the Purine-ring (Fig. 2c) and altered dynamics in the nucleotide pocket in a manner that might affect ADP release.

**Figure 1.**
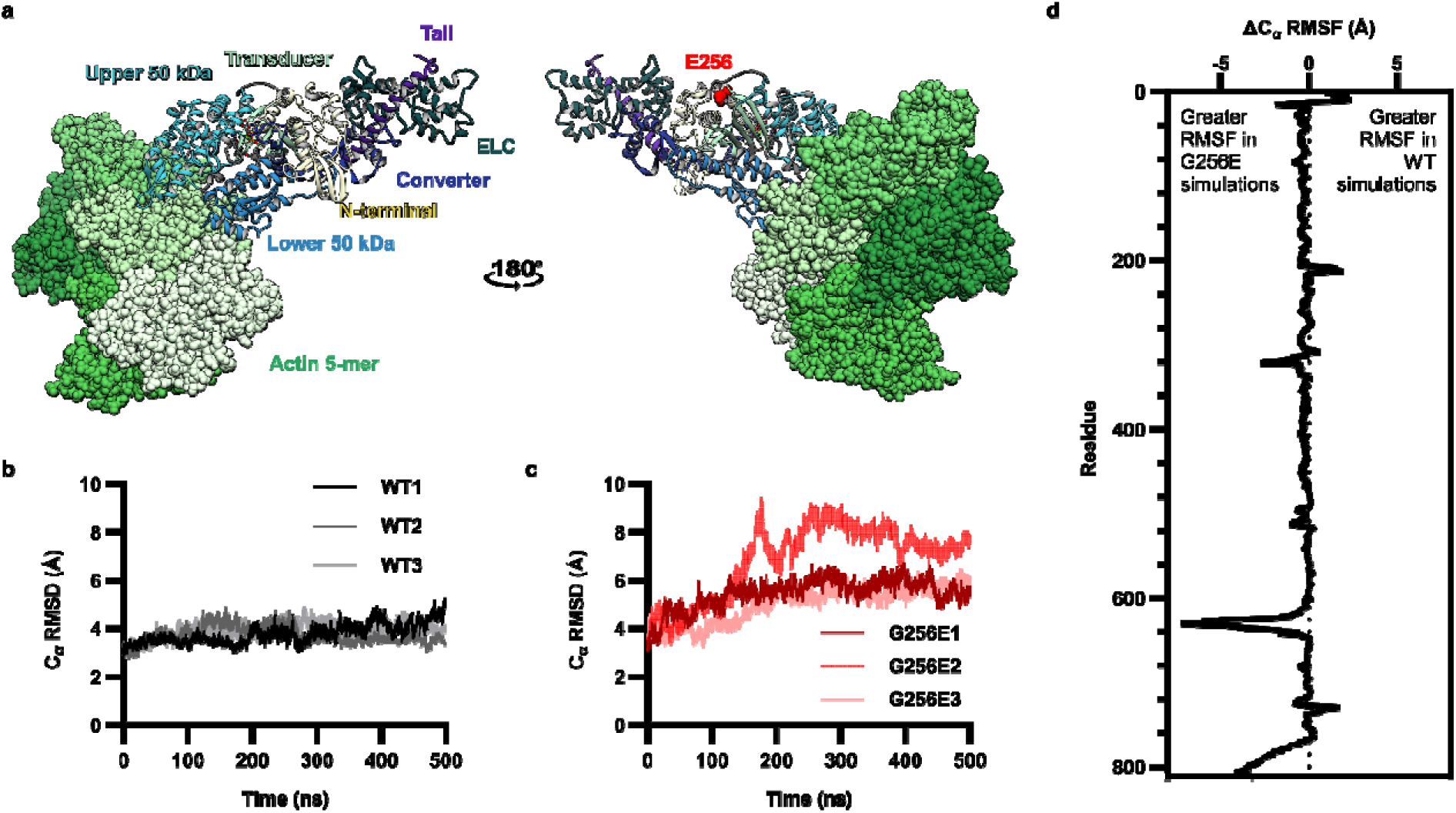
(a) The post-powerstroke model includes 5 actin monomers (light green, space-filling atoms), myosin motor domain, and essential light chain (ELC, dark green). The functional regions of myosin are colored uniquely: upper 50 kDa domain (cyan), lower 50 kDa domain (blue), transducer (seafoam green), N-terminal domain (yellow), converter domain (dark blue), and tail (purple). E256 (red) is located in a β-hairpin turn between beta strands β_6_ and β_7_ in the transducer region. (b) Global C_α_ RMSD of WT simulations and (c) G256E simulations. On average, G256E simulations had higher global C_α_ RMSD than WT simulations. (d) C_α_ RMSF difference between WT and G256E simulations by residue. Positive values indicate higher flexibility in WT simulations whereas negative values indicate higher flexibility in G256E simulations. C_α_ RMSF mapped onto WT and G256E structures can be visualized in Extended Data Fig. 1.

**Figure 2.**
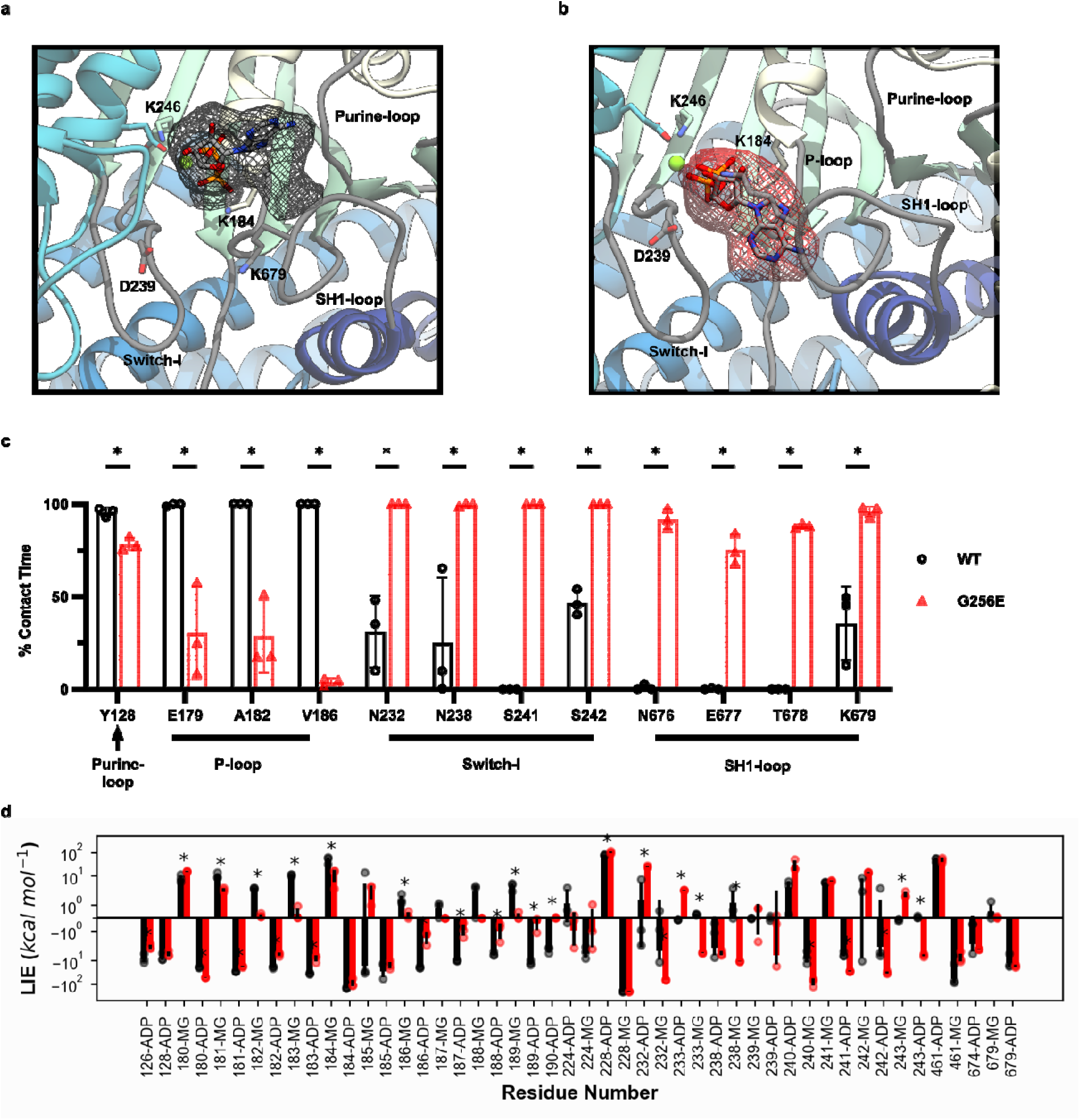
ADP occupancy maps for (a) WT and (b) G256E simulations show that ADP adopts a novel conformation in G256E simulations. Occupancy maps cover two standard deviations from the mean ADP occupancy. (c) Changes in ADP contacts with nucleotide pocket residues. In G256E simulations, contacts with the purine-loop and P-loop are lost while new contacts with Switch-1 and the SH-loop are formed. (d) Linear interaction energy analysis for the WT (black) and G256E (red) conventional MD simulations. The bars correspond to the average interaction energy between ADP or MG and myosin residues. Differences with p <= .05 are denoted by *.

The conformation of ADP within the nucleotide binding pocket was assessed by calculation of 6 distinct dihedral angles within ADP: θ_1_ (O1B-PB-O3A-PA), θ_2_ (PB-O3A-PA-O5′), θ_3_ (O3A-PA-O5′-C5′), θ_4_ (PA-O5′-C5′-C4′), θ_5_ (O5′-C5′-C4′-O4′), and θ_6_ (O4′-C1′-N9-C8). The dihedral angles were analyzed via 2D Ramachandran-style plots (Extended Data Fig. 1d-f). The quantified conformational entropy (Table 1) and percent coverage (Table 2) indicate that ADP is less flexible in the pocket and adopts a more stable conformation for G256E compared to WT simulations. We calculated the interaction energies between myosin and ADP.Mg^2+.^ Residues with at least 3 kT of interaction energy were considered part of the nucleotide binding pocket (residues 126, 128, 180-190, 224, 228, 232, 233, 238-243, 246, 461, 674, and 679) and most nucleotide binding pocket residues had statistically different interaction energies with ADP.Mg^2+^ (Fig. 2d). We also observed a statistically different center of mass distance between ADP.Mg^2+^ and the nucleotide binding pocket residues for G256E. The altered conformation of ADP.Mg^2+^ in the nucleotide binding pocket led to statistically distinct interaction energies with myosin residues (Fig. 2d).

**Table 1.** Conformational entropy of ADP in WT and G256E simulations.

| Dihedral pair | ADP entropy (cal mol<sup>-1</sup> K<sup>-1</sup>) |  |
| --- | --- | --- |
|  | WT | G256E |
| $\theta_1$ vs. $\theta_2$ | 9.6284 | 7.623 |
| $\theta_3$ vs. $\theta_4$ | 10.0047 | 8.9728 |
| $\theta_5$ vs. $\theta_6$ | 11.2456 | 10.2952 |

**Table 2.** Percent coverage of dihedral maps sampled by ADP in WT and G256E simulations.

| Dihedral pair | % coverage |  |
| --- | --- | --- |
|  | WT | G256E |
| $\theta_1$ vs. $\theta_2$ | 8.93% | 3.88% |
| $\theta_3$ vs. $\theta_4$ | 21.06% | 11.21% |
| $\theta_5$ vs. $\theta_6$ | 33.49% | 17.48% |

### 4.2 ADP release from the post-powerstroke state required more work for G256E simulations compared to WT

We used steered molecular dynamics (sMD) simulations to simulate ADP release from the nucleotide binding pocket of myosin in complex with actin (AM.ADP*). To create a reaction coordinate for steering, we calculated the linear interaction energy (lie) between myosin residues and either ADP or Mg^2+^ in the cMD simulations. Myosin residues with an average lie of at least 3 kT with either ADP or Mg^2+^ were selected as nucleotide binding pocket residues in the reaction coordinate (Fig. 2d). Next, we calculated the modal distance between ADP.Mg^2+^ and the nucleotide binding pocket residues in the cMD simulations (WT: 3.6 Å; G256E: 4.4 Å). We then selected 120 structures from the cMD simulations (approximately 20 structures from each replicate) that had ADP.Mg^2+^ - nucleotide binding pocket residue center of mass distances within 0.1 Å of the model results. Then, steered MD simulations were performed (Fig. 3d,e). On average, G256E myosin required 1.5 times as much work (WT: 58 kcal mol^-1^ G256e: 89 kcal mol^-1^, p = 2.4 x 10^-18^) to displace ADP.Mg^2+^ from the binding pocket, indicating a stronger interaction between G256E myosin and ADP.Mg^2+^ (Fig. 3a). The Jarzynski averaged trajectories followed this trend (WT: 26 kcal mol^-1^ G256E: 68 kcal mol^-1^, a 1.6-fold difference) (Fig. 3b,c). Note that the free energy values along the reaction coordinate are dependent on the pulling speed in the sMD simulations, and relative differences in free energy along the profile are more meaningful than absolute energies here.^47^ The free energy profiles show an initial steep rise along the reaction coordinate that corresponds to movement of the α and β phosphates followed by a shallower rise. More work was required to displace ADP.Mg^2+^ from G256E myosin due to the burial of the adenine ring in the SH1-loop in addition to increased interactions of the phosphate groups with Switch-1. We calculated the average interactions that ADP.Mg^2+^ made as they were steered out of the nucleotide binding pocket (Extended Data Fig. 3a,b). We found that ADP.Mg^2+^ formed different interactions with WT and G256E myosin as it left the pocket, in large part due to the burial of the adenine ring in the SH1-loop and increased interactions of the phosphates with Switch-1. For both the WT and G256E simulations, ADP was displaced from the nucleotide binding pocket and at the end of the simulations there were no interactions between ADP.Mg^2+^ and P-loop or purine binding loop residues. However, at the end of every simulation ADP.Mg^2+^ was still interacting with at least E228. Attempts to steer ADP.Mg^2+^ further along the reaction coordinate resulted in local unfolding of myosin. We hypothesize this is due to strong interaction with E228 and a secondary reaction coordinate may be necessary to further steer ADP.Mg^2+^ from this residue and based on visual inspection we do not predict differences in ADP release from this point onward. Overall, these data suggest increased binding affinity of ADP.Mg^2+^ for G256E myosin facilitated through modified ADP coordination that results in an altered release trajectory.

**Figure 3.**
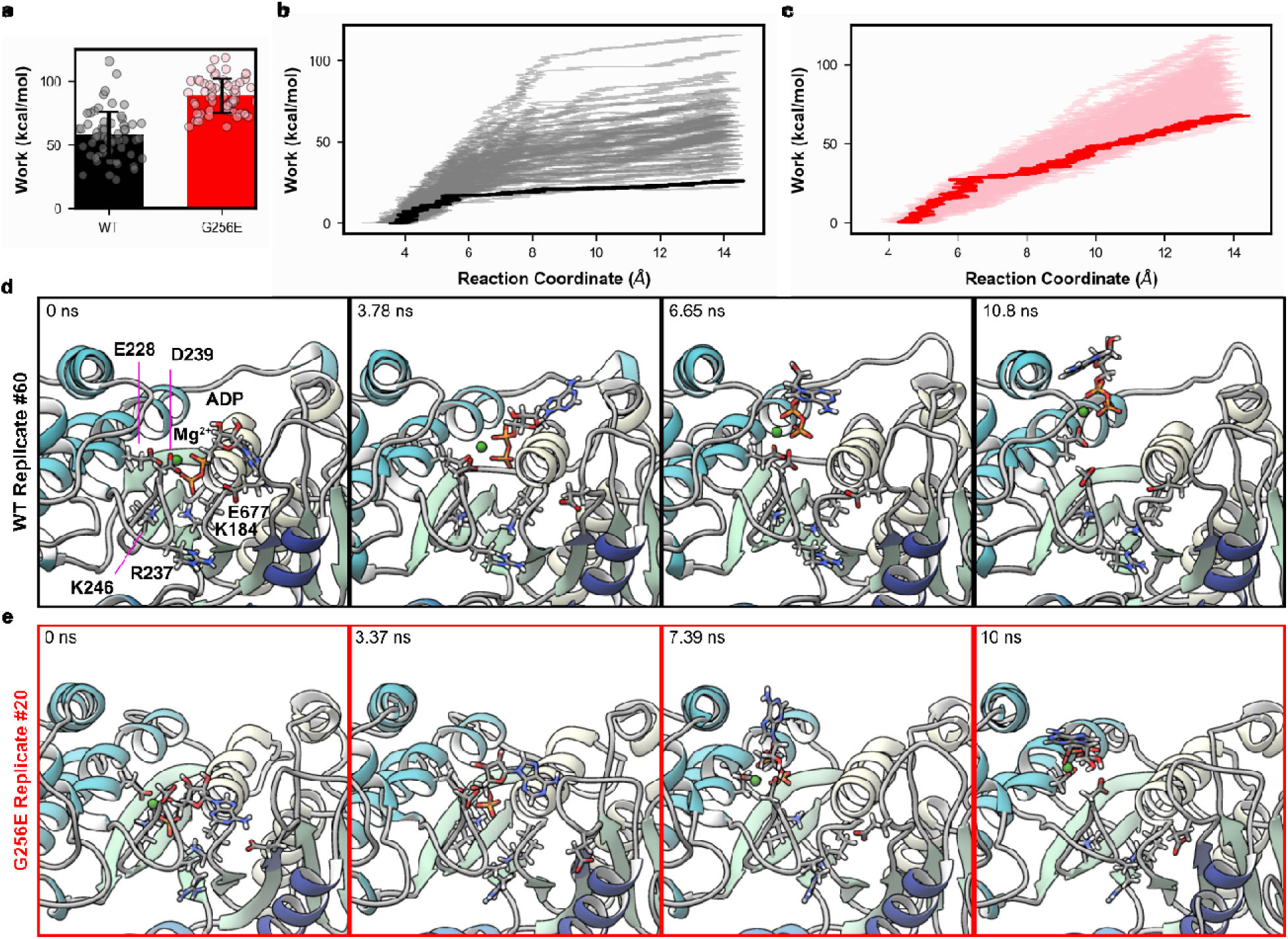
(a) Average work required to release ADP from WT (black) or G256E (red) myosin. On average G256E required 1.5x as much work to displace ADP. The free energy profiles of ADP release along our specified reaction coordinate are shown in panels (b) and (c) for the WT and G256E simulations. All 60 profiles are shown and the Jarzynski averaged profile is in bold. These plots show the energy required to move ADP a certain distance away from the nucleotide binding pocket. Snapshots that depict ADP release from myosin from the Jarzynski average trajectories are shown in panels (d) and (e) for the WT and G256E simulations, respectively.

**Figure 4.**
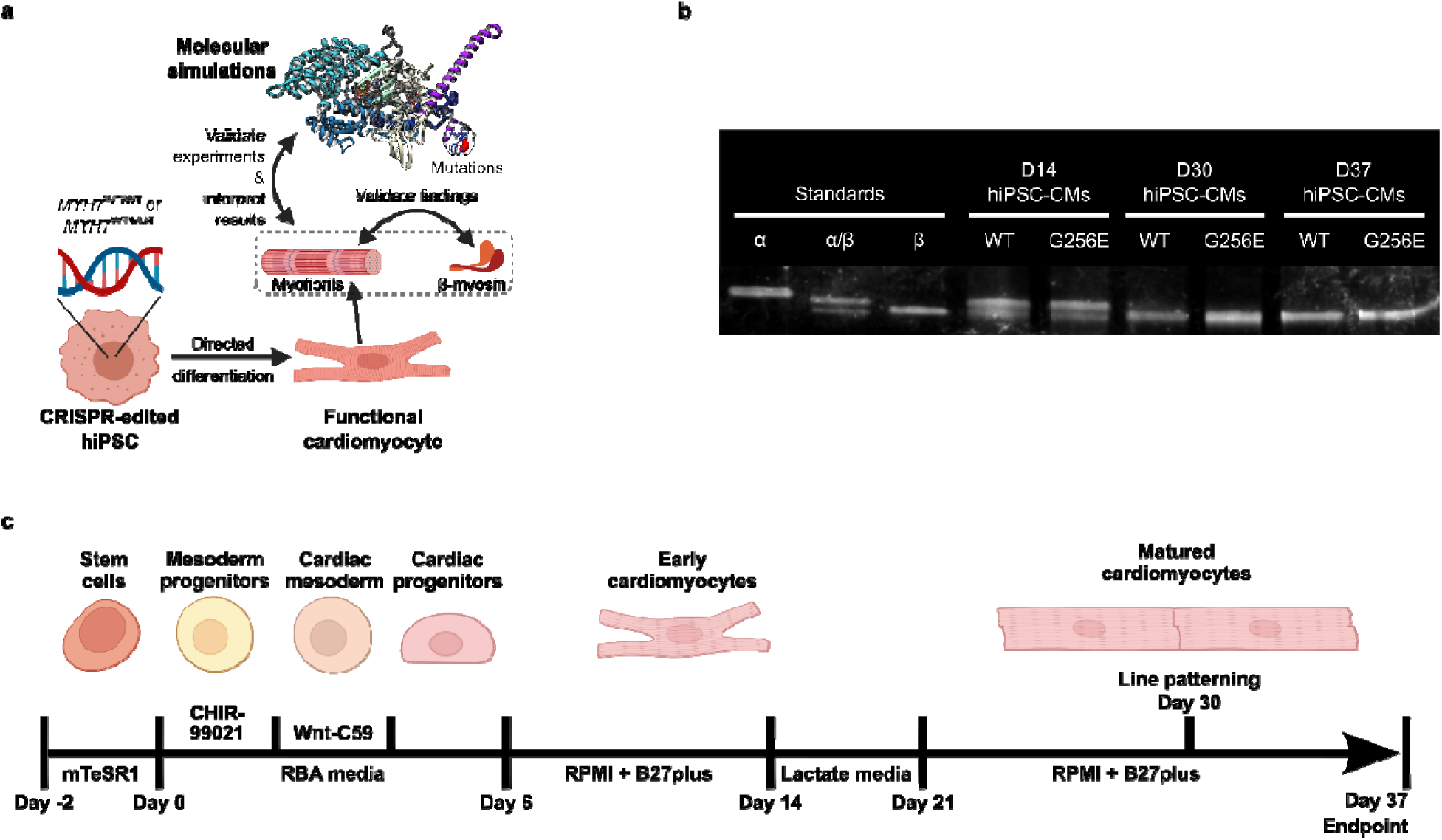
(a) Schematic of experimental and computational workflow synergy. Data from myofibrils derived from CRISPR-edited hiPSCs supported by biochemical experiments with purified β-myosin. Experimental findings are validated by molecular simulations. (b) Cardiac directed differentiation of hiPSCs. hiPSC-CMs are matured through lactate purification and line patterning for seven days before being skinned on D37 for myofibril mechanics. (c) SDS-PAGE myosin separation gel showing the conversion of α-MHC of D14 hiPSC-CMs to β-MHC by D30 and D37. Standards are from adult mouse atria (α-MHC), adult rat ventricle (α/β-MHC), and adult rabbit ventricle (β-MHC).

### 4.3 G256E sS1 has greater affinity for ADP

Next, to experimentally determine the ADP affinity of myosin (*K*_ADP_) we performed an ATP/ADP competitive binding assay using recombinant human β-cardiac short subfragment-1 (β-sS1), a single headed myosin construct containing only the human ventricular essential light chain. Experiments were performed by rapidly mixing 0.125 μM pyrene-actin.β-sS1 with 25 μM ATP + varying amounts of ADP (0, 20, 40, 80, 160, 320, and 640 μM) (all reported concentrations are post-mixing, Fig. 5a). In the absence of ADP, experimental curves were fit with a single-exponential which describes ATP-induced actomyosin dissociation. In the presence of ADP, experimental curves were fit with a double-exponential to account for the slow phase, which describes ADP dissociation from actomyosin. We plotted the relative *k*_obs_ of the fast phase (*k*_rel_ = *k*_obs_/ *k*_obs,0_) vs. ADP concentration to assess ADP affinity (Fig. 5b). G256E β-sS1 demonstrated ∼30% increased ADP affinity (*K*_ADP_ = 33 ± 3 µM, Fig. 5c) compared to WT β-sS1 (*K*_ADP_ = 48 ± 11 µM).

**Figure 5.**
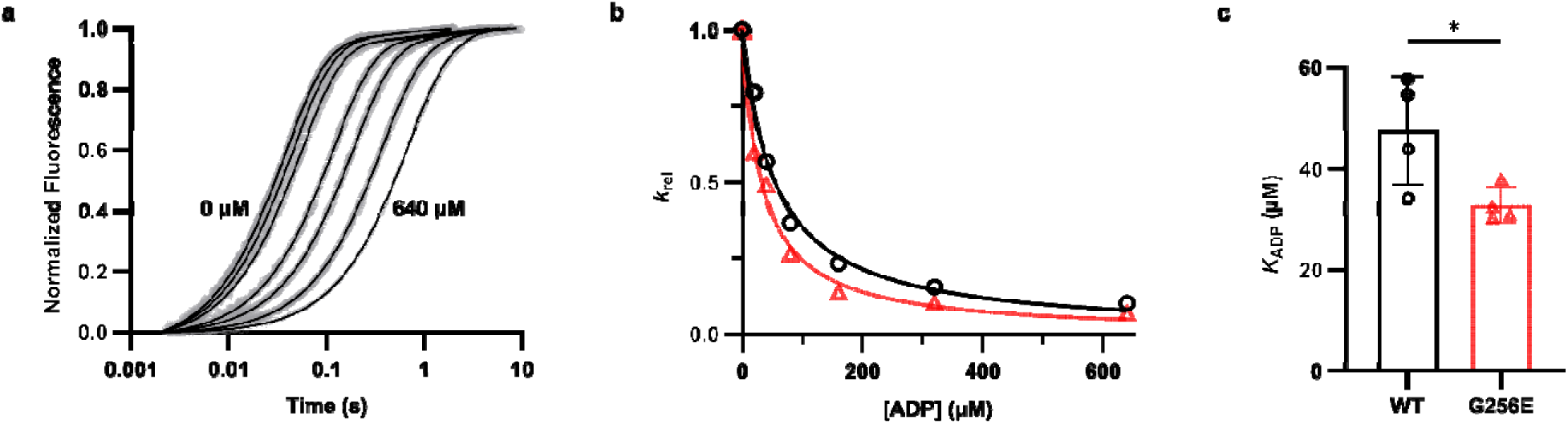
(a) Average traces (8-10 individual traces) of ATP-induced pyrene-actin.WT sS1 dissociation. Traces in the absence of ADP (0 µM) were fit with a single exponential. Traces in the presence of ADP (left to right: 20, 40, 80, 160, 320, 640 µM post-mixing) were fit with a double exponential. (b) The relative *k*_obs_ (*k*_rel_ = *k*_obs_/ *k*_0_ where *k*_0_ = *k*_obs_ when ADP = 0 µM) of the fast phase plotted vs. [ADP]. The data was fit with the equation *k*_rel_ = 1/(1 + [ADP]/*K*_ADP_) to obtain the apparent affinity. One representative trace each from WT and G256E are shown in (b). (c) G256E caused a ∼30% increase in ADP affinity (*K*_ADP_ = 32.82 ± 1.60 μM) compared to WT (*K*_ADP_ = 46.46 ± 3.23 μM). Error bars represent ± SD, n = 4.

### 4.4 G256E myofibril relaxation is slower and less sensitive to elevated ADP

To determine how the increased ADP binding affinity affects contractile properties, particularly the early, linear phase of relaxation that reflects crossbridge detachment, we used myofibrils from CRISPR/Cas9-edited hiPSC lines containing the heterozygous expression of the G256E mutation (*MYH7*^WT/G256E^) and an isogenic control (*MYH7*^WT/WT^). The cell lines underwent a series of quality control experiments to confirm genotype and validate that the gene-editing process did not have any off-target effects.^8^ We confirmed that our stem cell differentiation and culture conditions (Fig. 4b) provided hiPSC-CMs that express predominantly β-MHC by D30, validating the D37 timepoint was appropriate for experiments (Fig 4c). The mechanical and kinetic properties of hiPSC-CM myofibrils were measured in a custom-built apparatus using a fast solution switching (∼5 ms) technique, as previously described.^46^ Myofibrils were activated in a submaximal calcium solution (pCa 5.6), relaxed (pCa 9.0), then activated and relaxed with solutions containing 50% ADP (pCa 5.6ADP, 2.5 mM ATP + 2.5 mM ADP) to observe the effects of elevated ADP on contractile properties. Fig. 6a shows representative traces for contraction and relaxation and Fig. 6b shows a higher resolution view of the early, linear phase of relaxation, for measurements of the slope (*k*_REL,slow_) and duration (t_REL,slow_). Force generated at both pCa 5.6 and pCa 5.6ADP was significantly increased for G256E myofibrils by ∼2X (Fig. 6c, *p* < 0.001) compared to WT myofibrils. Activation kinetics (*k*_ACT_) with elevated ADP were decreased by ∼30% (Fig. 6d, *p* < 0.01) for both G256E and WT myofibrils. For relaxation, the main, fast phase rate (*k*_REL,fast_) was ∼30% slower for G256E myofibrils compared to WT (Fig. 6e, *p* < 0.05). Addition of ADP slowed *k*_REL,fast_ for both WT and G256E, but the magnitude of change was greater for WT *k*_REL,fast_, such that this rate was no longer significantly different for G256E vs WT myofibrils. For the early, linear phase of relaxation, the duration (t_REL,slow_) was prolonged approximately 2-fold by the addition of ADP for both G256E and WT myofibrils, as expected, but was not significantly different between G256E and WT myofibrils at pCa 5.6 or 5.6ADP. However, the slow phase of relaxation (*k*_REL,slow_) was significantly impaired for G256E vs WT myofibrils with ATP only solutions (Fig. 6f, *p* < 0.001) at pCa 5.6, similar to what we previously reported.^8^ Importantly, elevated ADP had a large effect on *k*_REL,slow_ in WT myofibrils (*p* < 0.0001), greatly slowing this rate, while it resulted in no appreciable change in *k*_REL,slow_ for G256E myofibrils. This suggests that ADP release is already significantly impaired in G256E myofibrils, such that the response to ADP product inhibition is blunted.

**Figure 6.**
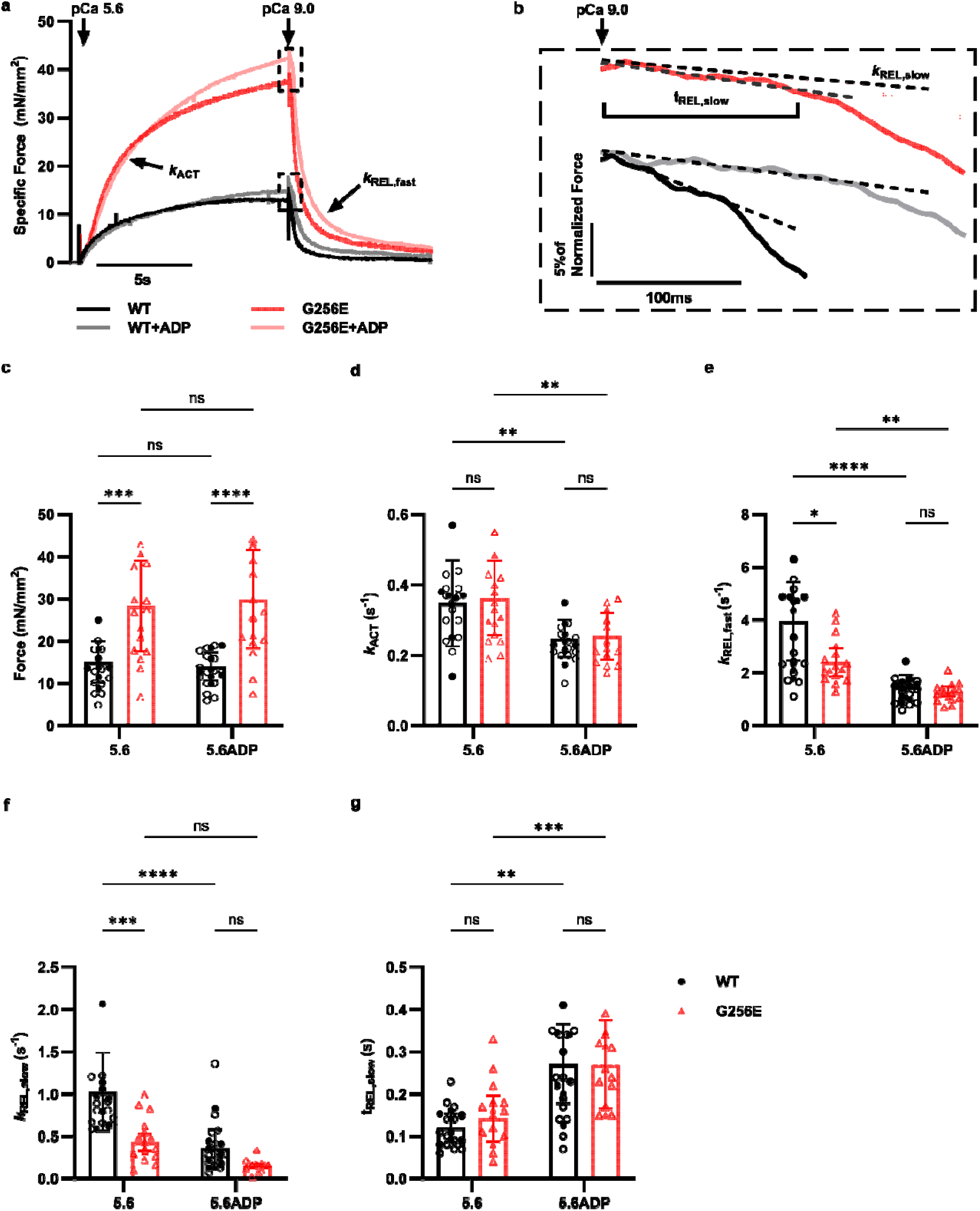
(a & b) Representative activation traces from hiPSC-CM *MYH7*^WT/G256E^ (red) and isogeni *MYH7*^WT/WT^ (black) myofibrils with a close-up of the slow, early phase of relaxation with linear fits (*k*_REL,slow_). Traces are an average of 2 sequential activations with activating solutions of 5.6 = pCa 5.6 activation paired with pCa 9.0 relaxation or 5.6ADP = pCa 5.6 activation + 50% ADP (2.5 mM ATP, 2.5 mM ADP) paired with pCa 9.0 + 50% ADP (2.5 mM ATP, 2.5 mM ADP) relaxation. (c) Max force generated is significantly higher in G256E myofibrils in both 5.6 and 5.6ADP activating solutions. ADP does not inhibit force generation. (d) Activation kinetics (*k*_ACT_) are unchanged between G256E and WT at both conditions (5.6 and 5.6ADP) and decreased with the addition of ADP. (e) G256E fast phase relaxation kinetics (*k*_REL,fast_) are slower at 5.6 compared to isogenic control; ADP inhibition decreases *k*_REL,fast_ significantly. (f) The kinetics of the slow, early phase (*k*_REL,slow_) are significantly decreased in G256E myofibrils at 5.6. The addition of ADP decreased *k*_REL,slow_ of WT myofibrils significantly, to the point at which they mimic the kinetics at G256E myofibrils at 5.6/5.6ADP. (g) The duration of the slow phase of relaxation (t_REL,slow_) is not significantly different when comparing G256E and WT. The addition of ADP prolongs the duration of this phase. Data are represented as mean ± SD. Statistical significance was determined by 2-way ANOVA. ns *p* > 0.05, \**p* < 0.05, \*\**p* < 0.01, \*\*\**p* < 0.001, \*\*\*\**p* < 0.0001, WT n = 15, G256E n = 13.

## 5. Discussion

### 5.1 Synergy between computation simulations and experiments

Here, we simulated ADP release from myosin using enhanced sampling computational techniques. We validated computational results with data collected from biochemical measurements of ADP affinity for sS1 myosin motors and mechanical measurements of hiPSC-CM-derived myofibrils. Our results highlight the synergy between computational simulation and experimental validation in understanding the structure-function basis of the myosin motor and contractile dysfunction resulting from disease associated mutations and this may provide intra-protein targets for more precise therapeutics approaches.

In post-powerstroke (AM.ADP) simulations, changing G256 to the E256 mutation forms an enduring salt bridge with R169. This was also observed in the M.ATP state simulations we reported recently.^8^ Even though this contact was consistent in simulations of both chemo-mechanical states, there were different consequences. In the M.ATP simulations, the salt bridge led to strain in sheets β5 and β6 of the transducer region and a reduction in the stability of hydrogen bonds between these sheets. However, in post-powerstroke (AM.ADP) simulations, the instability of the transducer region was not observed. This may be due to intrinsic differences in the nucleotide binding pocket in ‘closed’ (i.e. actin-free, ATP/ADP.P_i_-bound forms) vs ‘open’ (i.e. actin-bound, ADP-bound and nucleotide-free forms) conformations. These findings highlight the importance of simulating myosin in different chemo-mechanical states to obtain a more complete assessment of induced changes in structural dynamics, e.g. a single state structure analysis may result in incomplete understanding or incorrect conclusions. In the case of post-powerstroke simulations, it is possible that the presence of actin stabilized the transducer, as the transducer is known to adopt different conformations in the presence and absence of actin.^48,49^ The post-powerstroke simulations ultimately led to the discovery of altered nucleotide pocket dynamics due to the G256E mutation, which was consistent for all replicate simulations of the mutant. In particular, ADP formed novel contacts with the SH1-loop. This loop leads into the SH1 and SH2 helices, which link the converter domain to the rest of the myosin motor domain. In previous experimental studies, SH1-SH2 interactions were found to modulate the affinity of ADP to myosin, and a computational study found electrostatic interactions between the SH1 and SH2 helices alter ADP release.^50,51^ SH1 and the relay helix also connect the converter domain to the actin-binding surface, facilitating lever arm movement.^51^ Although we did not see any differences in either SH1-SH2 interaction or stability (i.e., secondary structure), it is possible some structural alterations could occur outside of our timescale (i.e., 500 ns).

Our sMD simulations represent the first computational study of ADP release from the actin-myosin complex, which was made possible by the existence of a high-resolution atomic structure of post-powerstroke myosin.^16^ The sMD simulations revealed that the conformation of ADP within the nucleotide pocket impacts ADP dissociation. In the case of G256E simulations, strong contacts between ADP and Switch-1 and the SH1-loop proved to be an extra energetic barrier inhibiting ADP release. sMD simulations also directed attention to particular residues that are important in ADP dissociation. K184, a positively charged residue in the P-loop lining the bottom of the pocket, is the first-line of defense in preventing ADP release, and the loss of a salt bridge between this residue and the β phosphate is necessary to initiate ADP release. Mutation of this residue to a glutamine (K184Q) has been seen in patients with HCM or left ventricular noncompaction cardiomyopathy and is currently a variant of unknown significance (VUS).^52^ Future studies of mutations in this residue and how it relates to ADP release may give insight into its potential pathogenicity. The next residue identified on the reaction coordinate is D239. In WT simulations, this residue forms a salt bridge with K679, acting as an ‘atomic-level staple’ to keep the nucleotide pocket closed. This residue is of particular interest because it has been tied to multiple disease-related mutations. D239N is a likely pathogenic mutation implicated in early-onset HCM.^53,54^ Based on our observations we hypothesize that the change from Asp, a negatively charged side chain, to the uncharged polar residue Asn, will disrupt the salt bridge between 239 and 679. This would likely result in faster ADP release. This may explain results from a study demonstrating faster actin gliding velocities and actin-activated ATPase rates in D239N myosin, as ADP release is the rate-limiting step in unloaded actin-myosin assays.^55^ Finally, K246 is another residue that is important for P_β_ stabilization. The mutation K246I is currently classified as likely pathogenic by Walsh et al.^56^ A change from positive residue to a non-polar one would likely affect coordination of the nucleotide in the pocket through destabilization of P_β_.^56^

In our previous study we found that, for isolated myofibrils from hiPSC-CMs, the kinetics of the slow, early phase of relaxation, *k*_REL,slow_ following maximal (pCa 4.0) calcium activation was reduced.^8^ This could result from either slowed ADP release from myosin or slowed ATP binding to myosin, as both steps that are required for cross-bridge detachment. Here we utilized elevated ADP as a product inhibition method to probe if ADP release from myosin is specifically altered with the G256E mutation. We chose to perform these experiments at a sub-maximal calcium level to make conditions more physiologically relevant, as pCa 5.6 is much closer to the rise in intracellular calcium that occurs during a cardiac twitch. Importantly, *k*_REL,slow_ was significantly slower for G256E myofibrils at pCa 5.6 (Figure 6F), a similar result as in our previous study at pCa 4.0.^8^ The addition of 2.5 mM ADP to WT myofibrils slowed *k*_REL,slow_ to a rate that was similar to that for G256E myofibrils in the absence of ADP, and addition of ADP for G256E myofibrils resulted in a significant but smaller further reduction of *k*_REL,slow_. The addition of ADP should inhibit ADP release and prolong the kinetics of this phase, which is seen in WT myofibrils but not G256E myofibrils. This suggests that the release of ADP is significantly compromised within G256E myofibrils, explaining in minimal response to ADP product inhibition. These data agree with the conclusions from our computational simulations and myosin biochemical measures regarding ADP.Mg^2+^ release and greater ADP.Mg^2+^ for myosin, respectively. However, we should not rule out other possible contributions to the relaxation rate, e.g. the rates of ATP binding and acto-myosin dissociation. Our preliminary stopped-flow measures of mant-ATP binding to S1 or HMM indicate that the rate of ATP binding is multi-fold faster than *k*_REL,slow_ for both WT and G256E myofibrils, suggesting ADP release may control myosin detachment and relaxation kinetics. This is supported by measurements of *k*_REL,slow,_ which was decreased for G256E compared to WT myofibrils at pCa 5.6 (Fig. 6f), and decreased by elevation of ADP for WT, but much less for G256E myofibrils.

### 5.2 A note on force development

At pCa 5.6, steady state force (Fig. 6c) was approximately doubled for G256E myofibrils compared with WT myofibrils, indicated increased calcium sensitivity of force, while activation kinetics, *k*_ACT_ (Fig. 6d) were not affected. This contrasts with measurements we reported for maximal activation (pCa 4.0) where both force and *k*_ACT_ were approximately doubled for G256E myofibrils.^8^ This suggests that the rate limiting chemo-mechanical process for the rate of force development at the sub-maximal calcium used (pCa 5.6) is new strong crossbridge formation and the force generating isomerization of myosin associated with inorganic phosphate release, and not ADP release or crossbridge detachment. The elevation of ADP did not affect steady state force but decreased *k*_ACT_ by a similar amount for both WT and G256E myofibrils. Together this suggests the mutation affects the number of myosins that bind, but not the binding and force development kinetics.

### 5.3 Limitations of our study

Here, we report the first all-atom MD simulations of β-cardiac myosin in the post-powerstroke state. However, our model includes only an actin pentamer and myosin residues 1-810. Thus, there is no effective load in our system and effects on ADP release related to tension (force) or interaction between more complex constituents of the sarcomere are not included. Some studies suggest Mg^2+^ may release from myosin prior to ADP release; however, here we have simulated release of ADP.Mg^2+^ as a united molecule. We have not yet simulated the rigor actomyosin complex for this mutation, nor have we explored differences in ATP binding or actomyosin attachment for G256E myosin. Additional impacts of thin filament regulatory proteins were also not considered in our current simulation studies. Finally, our models do not account for additional, more complex behavior of myosin, which includes destabilization of the interacting heads motif (IHM) that occurs with myofibril contractile activation and contributes to the hypercontractility of the G256E mutation.^8^ Stopped-flow biochemical experiments involve isolated actomyosin constructs in the absence of load which may over-report the effects the G256E mutation on ADP affinity, as load reduces the rate of ADP release from myosin.^57,58^ Additionally, shortened myosin constructs are unable to form the IHM found in native thick filaments. As such, we are unable to assess the combined effects or interplay between alterations of myosin recruitment (ON and OFF states) and crossbridge cycling. Myofibrils derived from hiPSC-CMs may contain fetal isoforms of sarcomeric proteins; however, our maturation protocol results in myofibrils that contain predominantly β-myosin, which is the adult ventricular isoform affected by the G256E mutation, and the crossbridge cycling rate is dominated by myosin.

To mitigate the limitations of any one technique, our interdisciplinary research strategy used different models to explore the effects of a mutation at different scales to identify the common feature (ADP release) that G256E affects.

## 6. Conclusions

In summary, we have demonstrated that the *MYH7* G256E impairs ADP release as well as the previously reported increase in myosin recruitment from the OFF state.^8^ Together, these changes contribute to the clinically observed HCM phenotype of hypercontractility and impaired relaxation through delayed cross-bridge detachment and a greater population of myosin in force bearing states. Our conventional MD results suggest that mutations in the transducer region can allosterically affect the nucleotide binding pocket. Using sMD, we developed a predictive method to computationally assess a specific transition step in the cross-bridge cycle–ADP release. Our results were corroborated with experimental data obtained from myofibrils obtained from hiPSC-CMs and recombinant sS1 protein, emphasizing the necessity for both computational approaches and validation against experimental data. Similar studies could help elucidate the mechanisms of other specific mutations in myosin implicated in disease and contribute to developing additional sarcomere targeted therapies.

## 7. Acknowledgements

This work used the Extreme Science and Engineering Discovery Environment (XSEDE) resource COMET through allocation TG-MCB200100 to M.R. XSEDE was supported by the National Science Foundation grant no. ACI-1548562. Partial funding for MCC was provided by award nos. T32HL007828 and K99HL173646 from the National Heart, Lung, and Blood Institute. Partial support for KYK was provided by T32EB032787. This research was supported by the University of Washington Center for Translational Muscle Research (CTMR) via the National Institute of Arthritis and Musculoskeletal and Skin Diseases of the National Institutes of Health award no. P30AR074990. This work was initiated and supported by the NIH/NIGMS grant 1RM1 GM131981-03 to JAS, KMR and MR and supported by NHLBI R01HL128368 to MR. This work would not have been possible without the Allen Institute for Cell Science team who contributed with cell line generation. The parental WT unedited hiPSC line, WTC, was provided by the Bruce R. Conklin Laboratory at the Gladstone Institutes and UCSF. The Allen Institute for Cell Science wishes to thank the Allen Institute for Cell Science founder, Paul G. Allen, for his vision, encouragement, and support. The content is solely the responsibility of the authors and does not necessarily represent the official view of the NHLBI or the NIH. The authors have no financial conflicts of interest that may be construed to these data.

## 8. Author Contributions

Designed research, M.C.C. and M.R.

Provided purified protein, D.P. and R.R.G.

Performed research, M.C.C., K.K., and T.S.M.

Analyzed data, M.C.C. and K.K.

Manuscript writing, K.K., M.C.C.,

M.R. Resources and Funding, M.R. and J.A.S.

## 11. Extended data

**Extended Figure 1.**
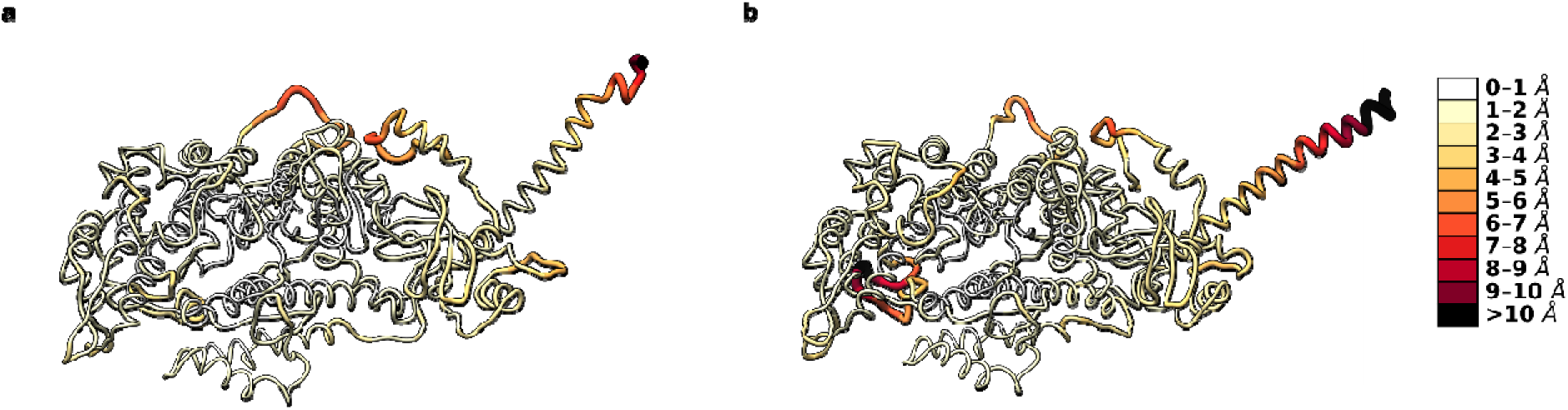
C_α_ RMSF mapped onto (a) WT and (b) G256E structures. Darker colors and thicker ribbons indicate higher C_α_ RMSF.

**Extended Figure 2.**
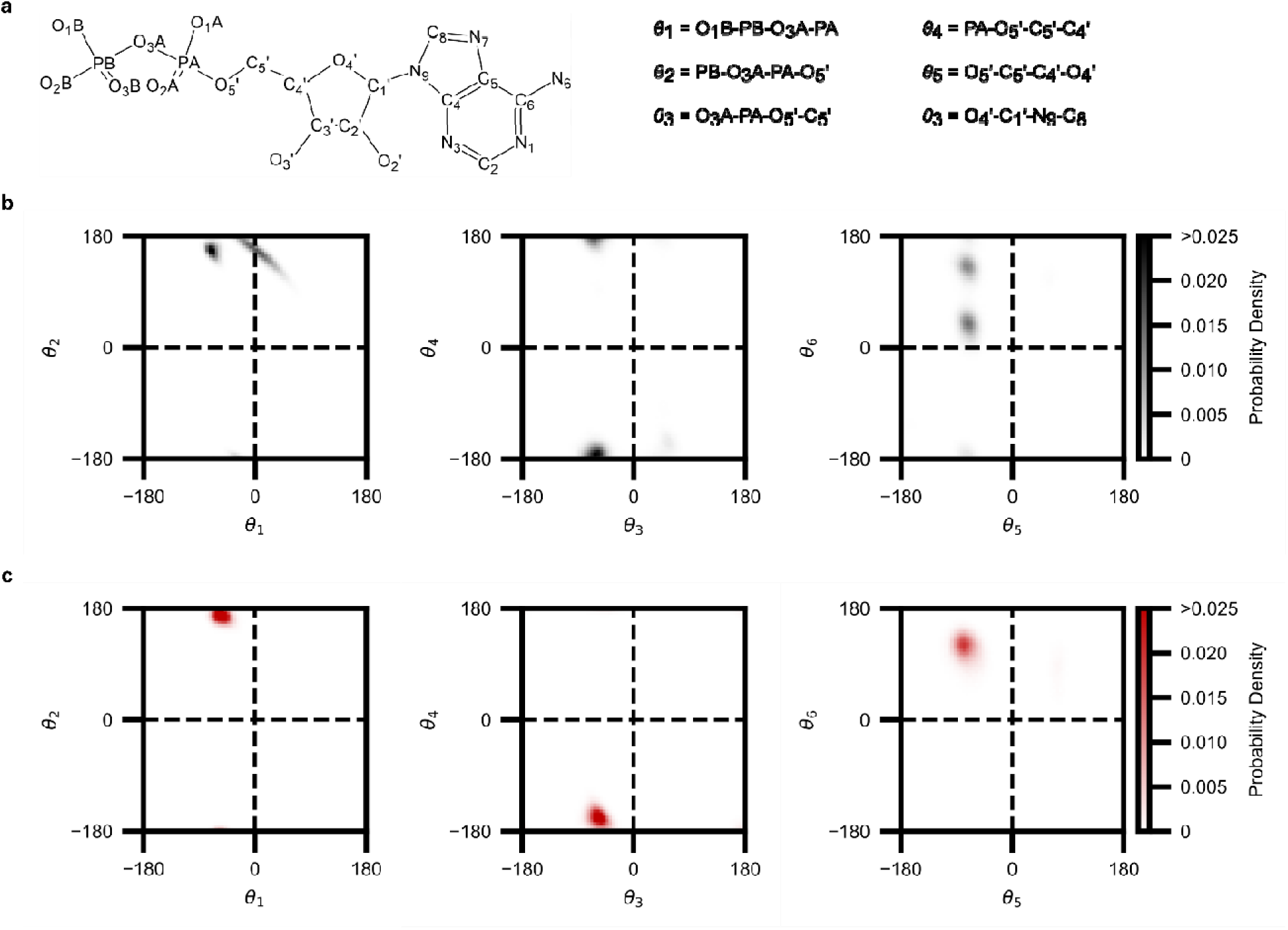
(a) ADP dihedral angle sampling for (b) WT and (c) G256E simulations. Six dihedral angles were defined: θ_1_ (O1B-PB-O3A-PA), θ_2_ (PB-O3A-PA-O5′), θ_3_ (O3A-PA-O5′-C5′), θ_4_ (PA-O5′-C5′-C4′), θ_5_ (O5′-C5′-C4′-O4′), and θ_6_ (O4′-C1′-N9-C8). These angles were plotted in Ramachandran-style 2-D histogram plots (bin size 5°x5°) to visualize changes in ADP conformation. Each bin is colored according to probability density.

**Extended Figure 3.**
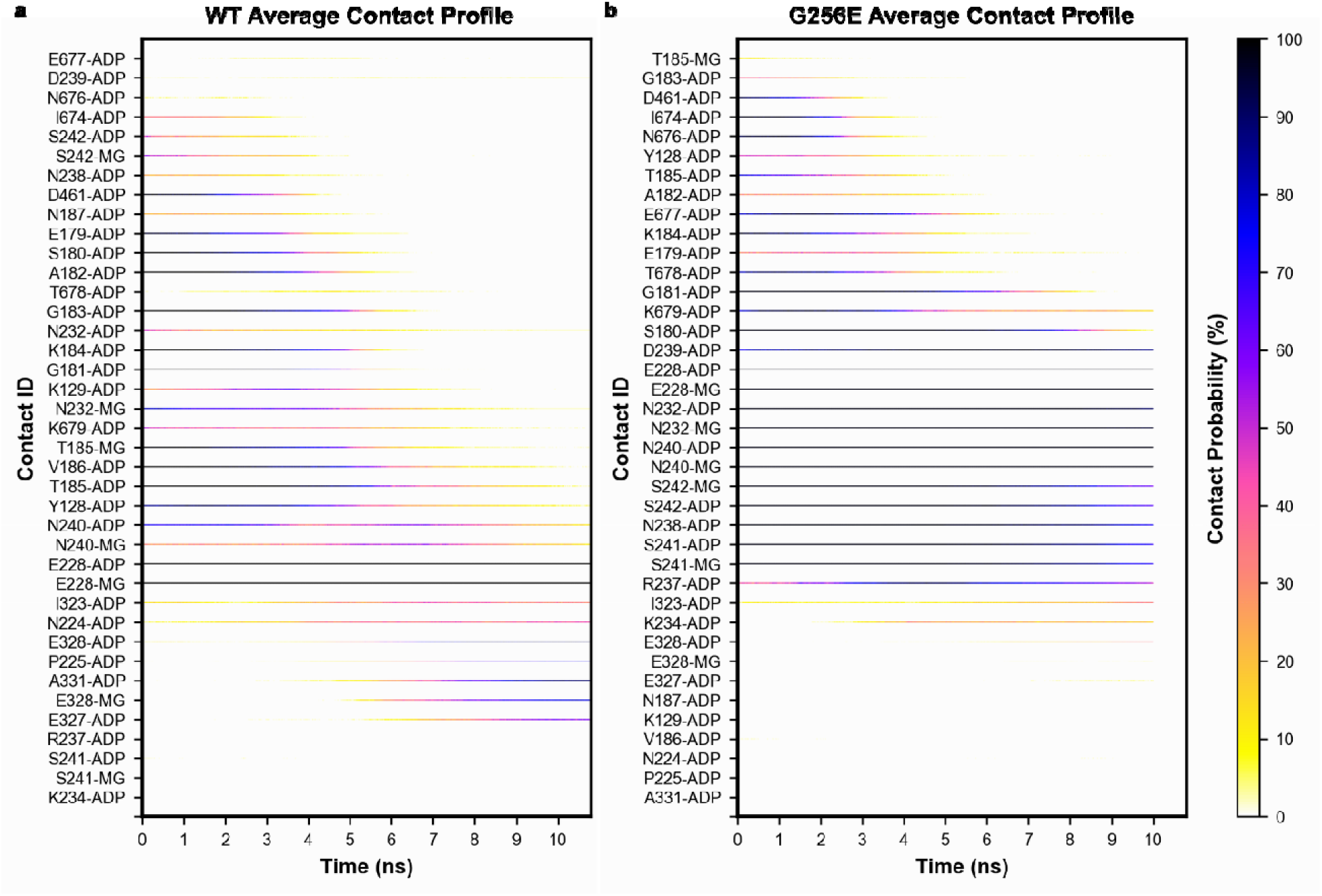
Panels (a) and (b) depict the average contacts made between myosin and ADP.Mg^2+^ during the steered MD simulations. Contacts are shown only if they appear at least once in at least 90% of the WT or 90% of the G256E simulations. For each contact (y-axis), we averaged the fraction of simulations for which the contact was present at each time in the SMD simulations -- this contact probability corresponds to the color of the dot plots. These plots show that ADP exits WT/G256E myosin via different pathways and highlights residues that form strong interactions with ADP during release (all black lines).

